# Timing of gene expression and recruitment in independent origins of CAM in the Agavoideae (Asparagaceae)

**DOI:** 10.1101/2021.11.10.468112

**Authors:** Karolina Heyduk, Edward V. McAssey, Jim Leebens-Mack

**Author notes:** **One sentence summary**: Independent origins of CAM in the Agavoideae show overall large similarity in diurnal gene expression profiles, but differential recruitment of the main CAM carboxylating enzyme. **List of author contributions**: K.H. conducted physiology experiments, sampled for RNA, prepared sequencing libraries, and analyzed data; E.V.M conducted molecular evolution analyses; J.L-M. helped with framing and data interpretation; all three authors contributed to writing and editing the manuscript.

## Abstract

CAM photosynthesis has evolved repeatedly across the plant tree of life, yet our understanding of the genetic convergence across independent origins remains hampered by the lack of comparative studies. CAM is furthermore thought to be closely linked to the circadian clock in order to achieve temporal separation of carboxylation and sugar production. Here, we explore gene expression profiles in eight species from the Agavoideae (Asparagaceae) encompassing three independent origins of CAM. Using comparative physiology and transcriptomics, we examined the variable modes of CAM in this subfamily and the changes in gene expression across time of day and between well-watered and drought-stressed treatments. We further assessed gene expression and molecular evolution of genes encoding phosphoenolpyruvate carboxylase (PPC), an enzyme required for primary carbon fixation in CAM. Most time-of-day expression profiles are largely conserved across all eight species and suggest that large perturbations to the central clock are not required for CAM evolution. In contrast, transcriptional response to drought is highly lineage specific. *Yucca* and *Beschorneria* have CAM-like expression of *PPC2*, a copy of *PPC* that has never been shown to be recruited for CAM in angiosperms, and evidence of positive selection in *PPC* genes implicates mutations that may have facilitated the recruitment for CAM function early in the evolutionary history of the Agavoideae. Together the physiological and transcriptomic comparison of closely related C_3_ and CAM species reveals similar gene expression profiles, with the notable exception of differential recruitment of carboxylase enzymes for CAM function.

## Introduction

The repeated origin of phenotypes across the tree of life has long fascinated biologists, particularly in cases where such phenotypes are assembled convergently - that is, using the same genetic building blocks. Documented examples of convergent evolution where the same genetic mechanisms are involved include the repeated origin of betalain pigmentation in the Caryophyllales (Sheehan et al., 2019), the parallel origin of caffeine biosynthesis in eudicots (Denoeud et al., 2014), and the repeated transition to red flowers in *Ipomoea* (Streisfeld and Rausher, 2009), among others. In all these cases, careful analysis of the genetic components underlying the repeated phenotypic evolution was driven by recruitment or loss of function of orthologous genes. Such convergence in the genetic mechanism suggests that the evolutionary path toward these phenotypes is relatively narrow, meaning the phenotype can only be obtained through a small set of very important molecular changes.

Such shared molecular mechanisms of repeated phenotypic evolution is especially surprising when observed across larger clades. For example, across all flowering plants, the large number of independent origins (~100) of both C_4_ and Crassulacean acid metabolism (CAM) photosynthesis imply relatively straightforward genetic and evolutionary paths from the ancestral C_3_ photosynthetic pathway (Edwards, 2019; Heyduk et al., 2019a). The overall photosynthetic metabolic pathway in C_4_ and CAM species is largely conserved; CO_2_ is converted to a four carbon acid by phosphoenolpyruvate carboxylase (PPC) and either moved to adjoining cells (C_4_) or stored in the vacuole overnight (CAM). The four carbon acids are then decarboxylated, resulting in high concentrations of CO_2_ in the cells where Rubisco is active. While some aspects of these photosynthetic pathways can vary among independent lineages, such as decarboxylation pathways in C_4_ lineages (Christin et al., 2009; Bräutigam et al., 2014), the same homolog of some genes has been repeatedly recruited for carbon concentration. In three independently derived C_4_ grass lineages, five out of seven photosynthetic genes examined had the same gene copy (orthologs) recruited, despite the presence of alternative copies (paralogs) of each gene (Christin et al., 2013). In *Cleome gynandra* (Cleomaceae) and *Zea mays* (Poaceae), transcription factors that induce expression of C_4_ photosynthetic genes in the required cell-specific manner were orthologous, despite >140 million years of evolution separating the two lineages (Aubry et al., 2014).

While C_4_ is known for the unique Kranz anatomy that allows the carbon concentrating mechanism to function efficiently, CAM instead relies on the temporal separation of CO_2_ assimilation and conversion of CO_2_ into sugars. The diurnal cycle of primary CO_2_ fixation and photosynthesis in CAM plants is thought to require a close integration with the circadian clock, though how that is explicitly accomplished remains unknown. Studies have shown only a handful of core clock genes differ in their expression between C_3_ and CAM species (Yang et al., 2017; Yin et al., 2018), though many of these studies rely on comparisons of distantly related species, confounding changes attributable to evolutionary distance with those that underlie the evolution of CAM photosynthesis. While the core clock seems largely similar in CAM and C_3_ species, 24-hour expression profiles for genes involved in carboxylation, decarboxylation, sugar metabolism, and stomatal movement have been shown to differ between C_3_ and CAM species (Ceusters et al., 2014; Ming et al., 2015; Abraham et al., 2016; Heyduk et al., 2018a; Wai et al., 2019), suggesting a regulatory link between clock genes and genes contributing to CAM function.

A hallmark of CAM is the evening expression of phosphoenolpyruvate carboxylase (*PPC*) genes, which produce the enzyme required for the initial fixation of atmospheric CO_2_ into an organic acid in both C_4_ and CAM plants. Unlike Rubisco, which has affinities for both CO_2_ and O_2_, PPC has only carboxylase function, which it uses to convert bicarbonate and phosphoenolpyruvate (PEP) into oxaloacetate (OAA). The carboxylating function of PPC is used by all plants to supplement intermediate metabolites into the tricarboxylic acid (TCA) cycle, and therefore *PPC* genes are present in all plant lineages in multiple copies. PPC enzymes employed by the CAM pathway are active in the evening and night, whereas TCA-related PPC enzymes are likely to have constitutive expression across the diel cycle, with perhaps higher activity during the day. Transcriptomic investigations of CAM species have shown that expression of the *PPC* genes involved in CAM is induced to much higher levels at dusk and overnight (Ming et al., 2015; Brilhaus et al., 2016; Yang et al., 2017; Heyduk et al., 2018a; Heyduk et al., 2019b): expression levels of CAM *PPC*s can be 100-1000x higher than *PPC* homologs contributing to housekeeping functions.

There are two main families of *PPC* genes in flowering plants: *PPC1*, which is typically present in 2-6 copies in most lineages (Deng et al., 2016), and *PPC2*, which shares homology with a *PPC* gene copy found in bacteria, and is typically found in single or low copy in plant genomes. *PPC1* forms a homotetramer, whereas *PPC2* requires the formation of a hetero-octamer with *PPC1* to function (O’Leary et al., 2009). *PPC1* is used in the TCA cycle in plants, and in all published cases within angiosperms a *PPC1* gene copy is recruited for CAM (and C_4_) function. *PPC2* has been shown to be involved in pollen maturation, fatty acid production in seeds, and possibly root development and salt sensing (Gennidakis et al., 2007; Igawa et al., 2010; Wang et al., 2012), though overall consensus on *PPC2* function in plants remains elusive.

To understand both the evolution of CAM, as well as the recruitment of *PPC* homologs in independent origins of CAM, we built upon existing physiological and transcriptomic data in the Agavoideae (Asparagaceae) by investigating additional species. CAM evolved three independent times in the Agavoideae: once in *Agave sensu lato* (*Agave s.l.,* includes the general *Agave, Manfreda, and Polianthes*), once in *Yucca*, and once in *Hesperaloe* (Heyduk et al., 2016b). Previous research compared gene expression and physiology in closely related C_3_ and CAM *Yucca* species (Heyduk et al., 2019b) and, separately, in species that range from weak CAM (low amounts of nocturnal CO_2_ uptake) to strong CAM in *Agave s.l. (Heyduk et al., 2018b)*. Gene expression profiles for key CAM genes in the C_3_ *Yucca* species studies showed CAM-like expression, especially when drought stressed, suggesting that perhaps *Yucca* or even the Agavoideae as a whole was primed for evolution of CAM due to gene regulatory networks and expression patterns that existed in a C_3_ ancestor. Here we conducted additional RNA sequencing in two species of *Hesperaloe* (CAM) and one species of *Hosta* (C_3_)) to assess 1) how gene expression varies in timing of expression and in response to drought stress across Agavoideae and 2) to what extent have the three independent origins of CAM in the Agavoideae involved recruitment of the same carboxylating enzyme gene homologs.

## Results

### CAM in the Agavoideae

Gas exchange and leaf titratable acidity amounts implicate CAM in both *Hesperaloe* species and C_3_ photosynthesis in *Hosta venusta* (Fig. 2, Supplemental Table S1 and Table S2). While we did not sample *Hosta venusta* under drought-stressed conditions, it’s thin leaf morphology (Heyduk et al., 2016b) and shady, mesic habitat suggests it is very unlikely to employ any mode of CAM photosynthesis. None of the CAM species in the Agavoideae examined here have high levels of nighttime CO_2_ uptake, and most still rely at least partially on daytime CO_2_ fixation by Rubisco (Fig. 2). Furthermore, *Yucca, Agave,* and *Manfreda* all appear to downregulate CAM under drought stress, as seen in both their gas exchange patterns and titratable acidity levels under drought relative to well-watered status (Fig. 2). *Hesperaloe parviflora* had slightly higher CO_2_ uptake at night than did *H. nocturna*, though both had appreciable levels of acid accumulation, and unlike *Yucca* and *Agave sensu lato* species, had a slight upregulation of CAM under drought stress. Finally, as previously described (Heyduk et al., 2018b), *Polianthes tuberosa* and *Beschorneria yuccoides* are C_3_+CAM, and both are able to facultatively employ CAM under drought stress.

**Figure 1.**
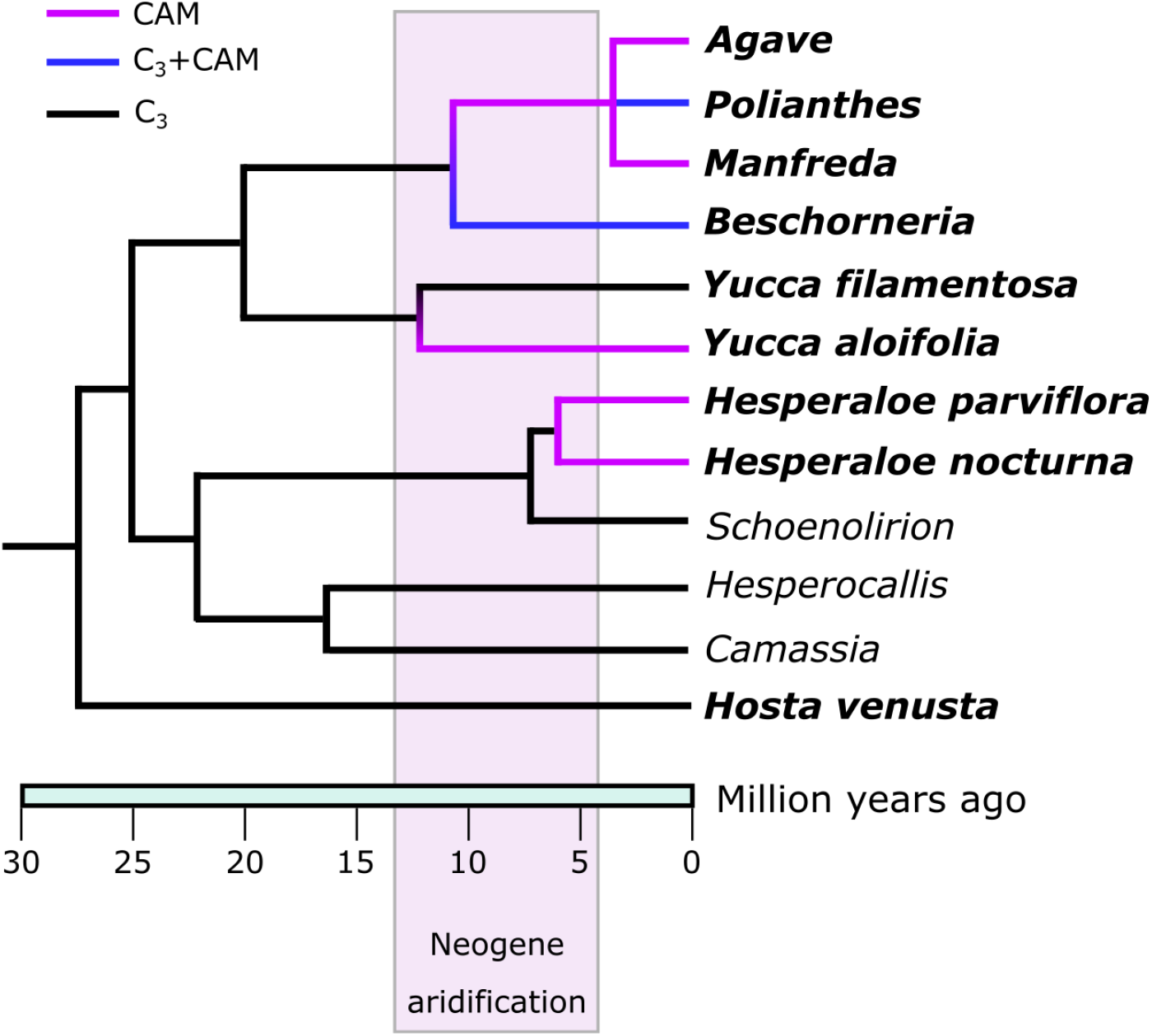
Simplified phylogeny of the Agavoideae, with estimated mean divergence times adapted from (McKain et al., 2016). Bolded taxa names are the species/genera included in this study. Branches are labeled according to photosynthetic pathway as described via previous work (Heyduk et al., 2016b; Heyduk et al., 2018b; Heyduk et al., 2019b).

**Figure 2.**
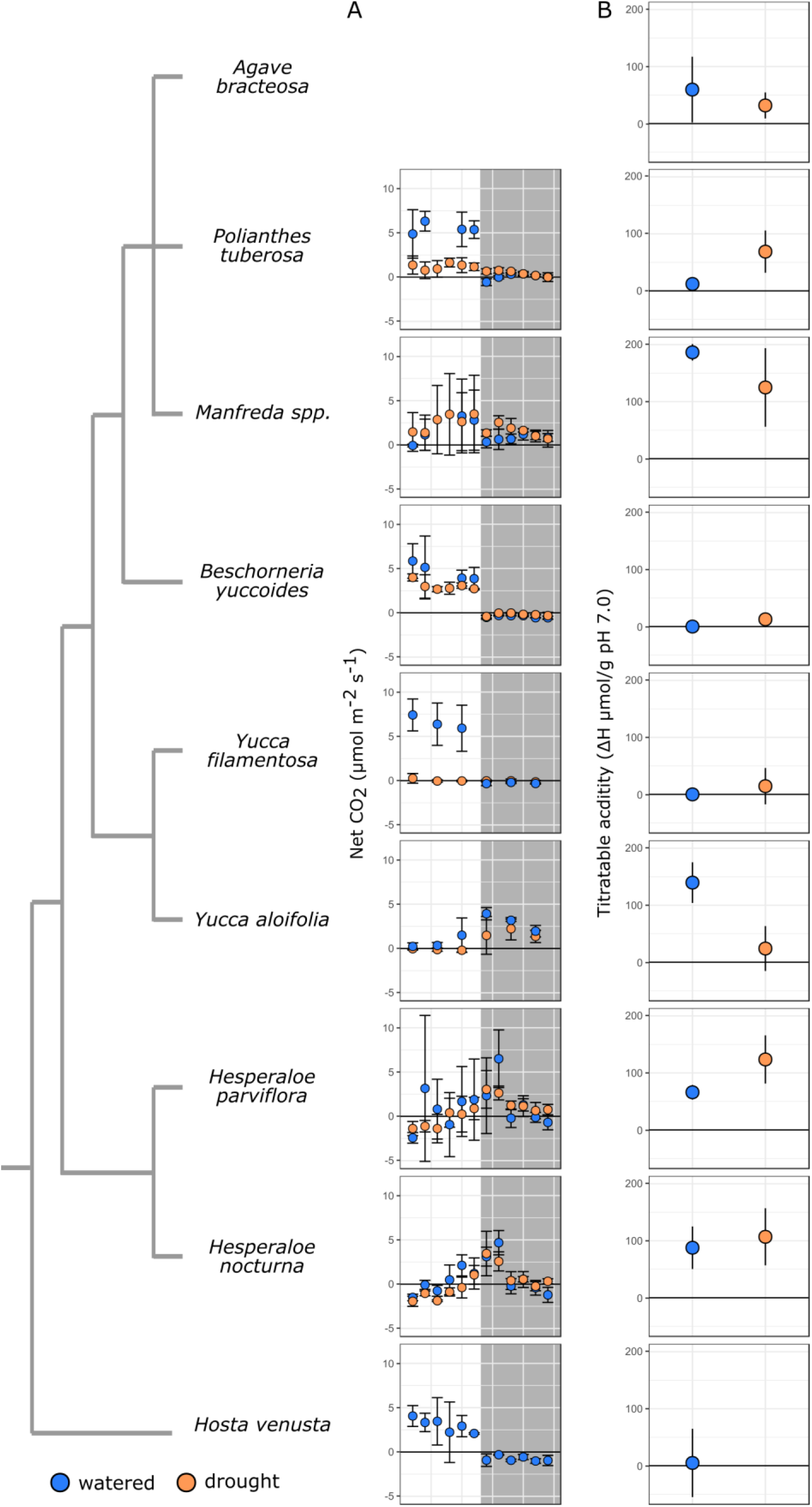
Photosynthetic physiology of species in the Agavoideae. Species relationships are represented by the cladogram to the left; gas exchange (A) and titratable leaf acidity (B) are shown per species. Data for all species except *Hesperaloe* and *Hosta* comes from previous work (Heyduk et al., 2018b; Heyduk et al., 2019b).

### Cross-Agavoideae comparisons

The number of transcripts that showed significant time-structured expression varied across species, with the fewest in the C_3_ species *Hosta venusta* (n=5,576) and the highest in the CAM species *Agave bracteosa* (n=28,856) (Fig. 3A). All species that use C_3_ photosynthesis or exhibit weak CAM had fewer transcripts that had significant change in diurnal expression, with the exception of *Polianthes tuberosa*, which uses CAM facultatively moreso than *Beschorneria yuccoides* does (Fig. 2). Many gene families (923) were time-structured in all 8 species (Supplemental Table S3); 731 additional gene families were time-structured in all species with the exception of *Hosta* (Fig. 3B, Supplemental Table S4). This latter set included a number of canonical CAM genes, including both *PPC1* and *PPC2*, as well as phosphoenolpyruvate carboxylase kinase (*PPCK*), a kinase dedicated to the phosphorylation of PPC, and thought to be required for efficient CAM (Taybi et al., 2000), auxin-related response genes, and a number of genes related to light reactions (e.g., photosystem II reaction center protein D).

**Figure 3.**
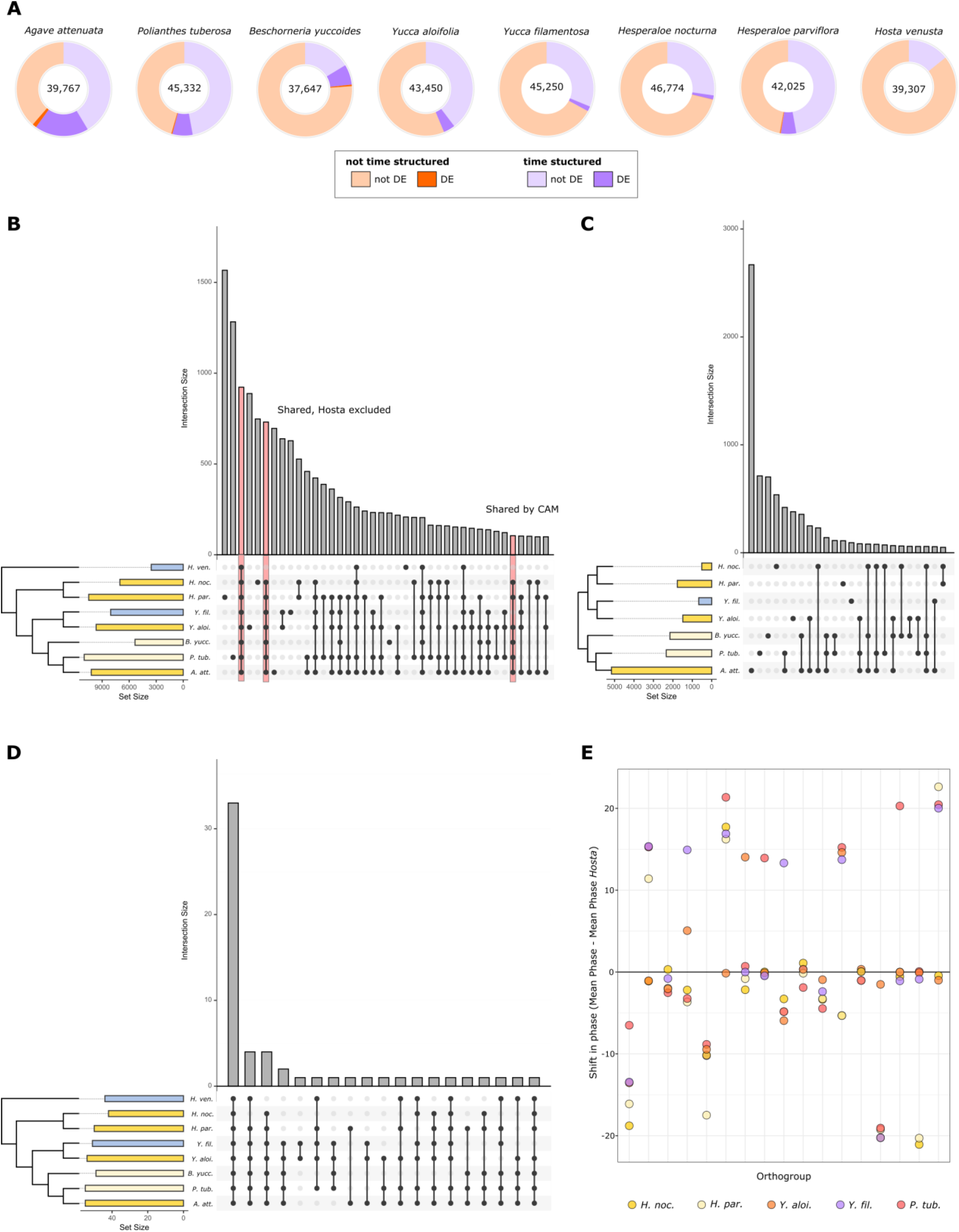
A) Total number of transcripts assessed in each species (center), with proportion of transcripts showing significant time-structured expression (orange shades). A subset of both time-structured and not time-structured transcripts were differentially expressed under drought conditions (purple shades). B) Upset plot showing overlap in gene families (orthogroups from OrthoFinder) that were time structured across the eight species; bars on left of species names indicate total number of gene families per species, colors indicate CAM (bright yellow), C_3_+CAM (pale yellow), and C_3_ (blue). C) Comparison of gene families that had differential expression under drought stress across seven of the eight species (*H. venusta* was not droughted). D) Comparison of gene families with core circadian clock annotations that had time-structured expression across all eight species. E) Shift in mean phase relative to *H. venusta* (mean phase of species - mean phase in *H. venusta*) in 17 gene families that had significantly different (p<0.01) cycling across species as indicated by Metacycle.

The number of genes responsive to drought stress was far lower than the total number with time-structured expression, and the majority of drought responsive genes had time-structured expression in at least one condition (watered or drought) (Fig. 3A,C). *Agave* had the largest number of differentially expressed genes under drought (~20%, Fig. 3A), while *H. nocturna* had the fewest (~2%). In both *Yucca* species, all drought-responsive genes were time-structured in their expression. Examination of shared gene families of drought-responsive genes across the species showed that many gene families were unique to a particular species (Fig. 3C), suggesting that drought response in the Agavoideae is variable and lineage-specific.

Of the gene families with circadian clock annotations, over half (33/58) had significant time-structured expression in all 8 species (Fig. 3D). Comparisons of phase (timing of peak expression) across the 8 species resulted in few differences in phase between species. In a comparison of CAM species (excluding *Agave* and *Beschorneria* due to low replicates/resolution; see methods) vs. C_3_ species, only four gene families had a significant shift in phase: *Pseudo-response regulator 9* (*PPR9*), *Alfin-like* (*AFL*), *telomere binding protein* (*TRFL*), and a gene of unknown function (no *Arabidopsis* homolog, and BLAST hits are uncharacterized proteins) (Supplemental Table S5). The comparison of phase changes between *Hosta* and the remainder of the Agavoideae species produced only a single gene family that had a shift in average timing of expression: *TRFL*, the same gene family found to be different between CAM and C_3_ species. In general, expression patterns across species were highly similar; of the 265 gene families that were 1) common in all 8 species and 2) significant cyclers as assessed by Metacycle, only 17 had a shift in phase when testing for species as an explanatory factor (p<0.01) (Fig. 3E)(Supplemental Table S6). In the majority of these gene families, the mean phase shift was low or were instances in which one species had a large phase shift different from the remaining species (Fig. 3E), but none had a concerted C_3_-to-CAM shift. In general, timing of expression was similar across all eight species in the majority of gene families.

### PPC expression

Gene tree reconstruction of sequences placed in *PPC1* and *PPC2* gene families by OrthoFinder are largely consistent with previous analyses (Fig. 4) (Deng et al., 2016; Heyduk et al., 2019a). The *PPC1* tree shows a duplication event within monocot evolutionary history, after the divergence of the Dioscorales (represented by *Dioscorea alata*) from the lineage leading to the last common ancestor of Asparagales and Poales (though *D. alata* appears to have a lineage-specific duplication). The monocot duplication event is independent from a similar duplication event in ancestral eudicots (Christin et al., 2014; Silvera et al., 2014). The placement of the *Acorus americanus* gene in the *PPC2* phylogeny as sister to all other sampled angiosperm homologs except *Amborella* is likely a result of the lack of other eudicots taxa in the analyses, or possibly eudicot-like mutations in the *A. americanus PPC2* gene. Regardless, the remainder of the gene tree is concordant with species relationships.

**Figure 4.**
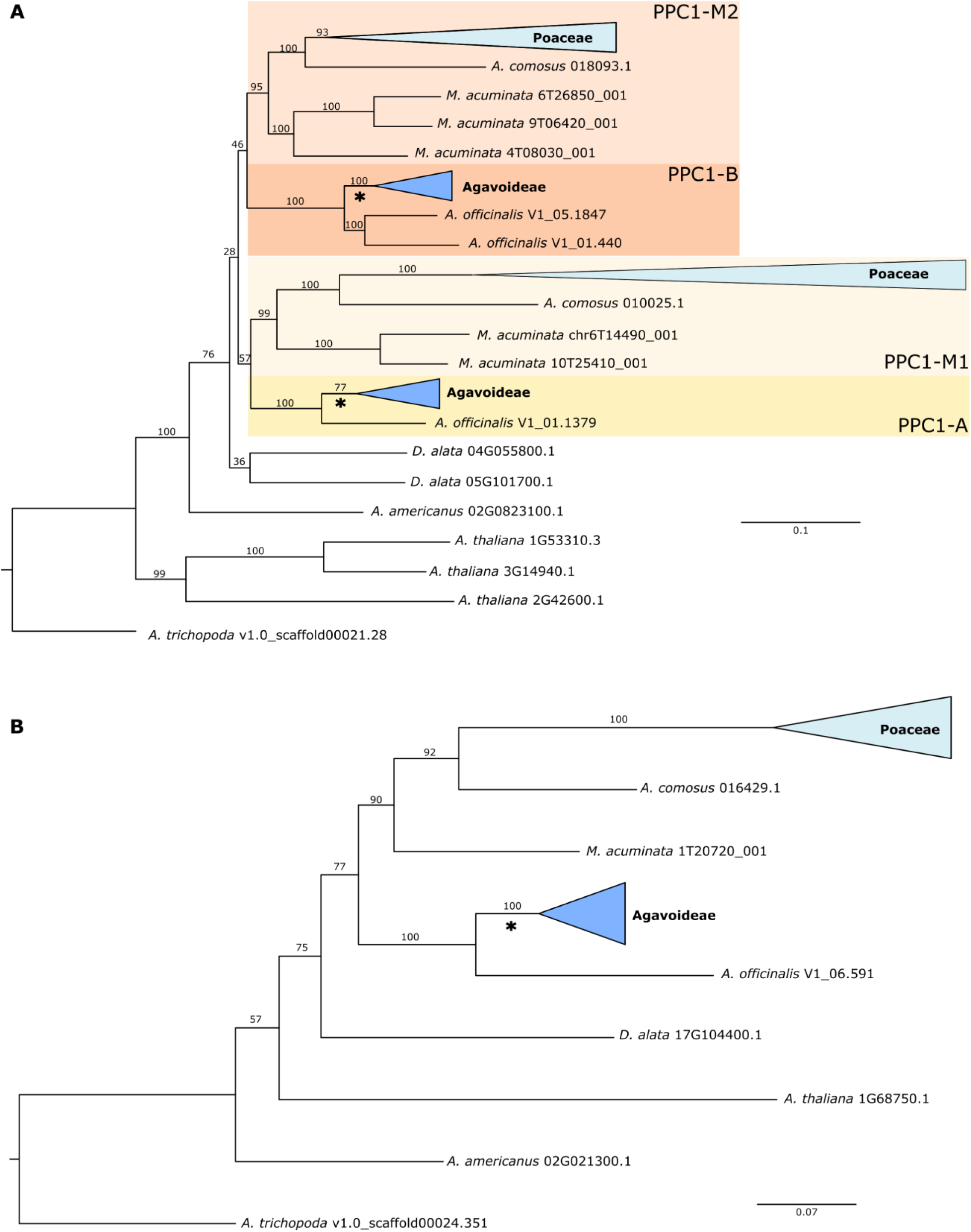
Gene trees estimated with IQTree for PPC1 (A) and PPC2 (B). Members of Poaceae and Agavoideae are collapsed for readability. All rapid bootstrap values are reported. Branches used for branch, branch x sites, and clade model tests in codeml are subtended by an asterisk.

While both major clades of *PPC1* were expressed in the Agavoideae, the overall expression levels of *PPC1-B* transcripts was much higher than *PPC1-A,* particularly in CAM species (Fig. 5). *PPC1-B* expression also increased under drought notably in *Polianthes tuberosa*, known to engage in facultative CAM upon drought stress (Fig. 2). *PPC2* transcripts were highly expressed in both *Yucca aloifolia* and *Beschorneria yuccoides,* strong CAM and facultative-CAM species, respectively (Fig. 6). Expression of *PPC2* increased with drought in *Beschorneria*, consistent with increased CAM activity under drought conditions (Fig. 2). Three gene copies of *PPC2* were identified in the *Yucca aloifolia* genome, and all three had characteristic CAM-like expression, with a peak before the onset of the dark period. Notably, *PPC2* is also expressed in a CAM-like pattern, albeit at lower levels, in the C_3_ *Yucca filamentosa* (Fig. 6). This finding is consistent with previous RNA seq analyses of *Yucca* (Heyduk et al., 2019b). *Hesperaloe nocturna* gene expression is not shown in Fig. 5 because the lengths of PPC transcripts were too short, and thus were filtered out from our gene tree estimation and subsequent expression analyses.

**Figure 5.**
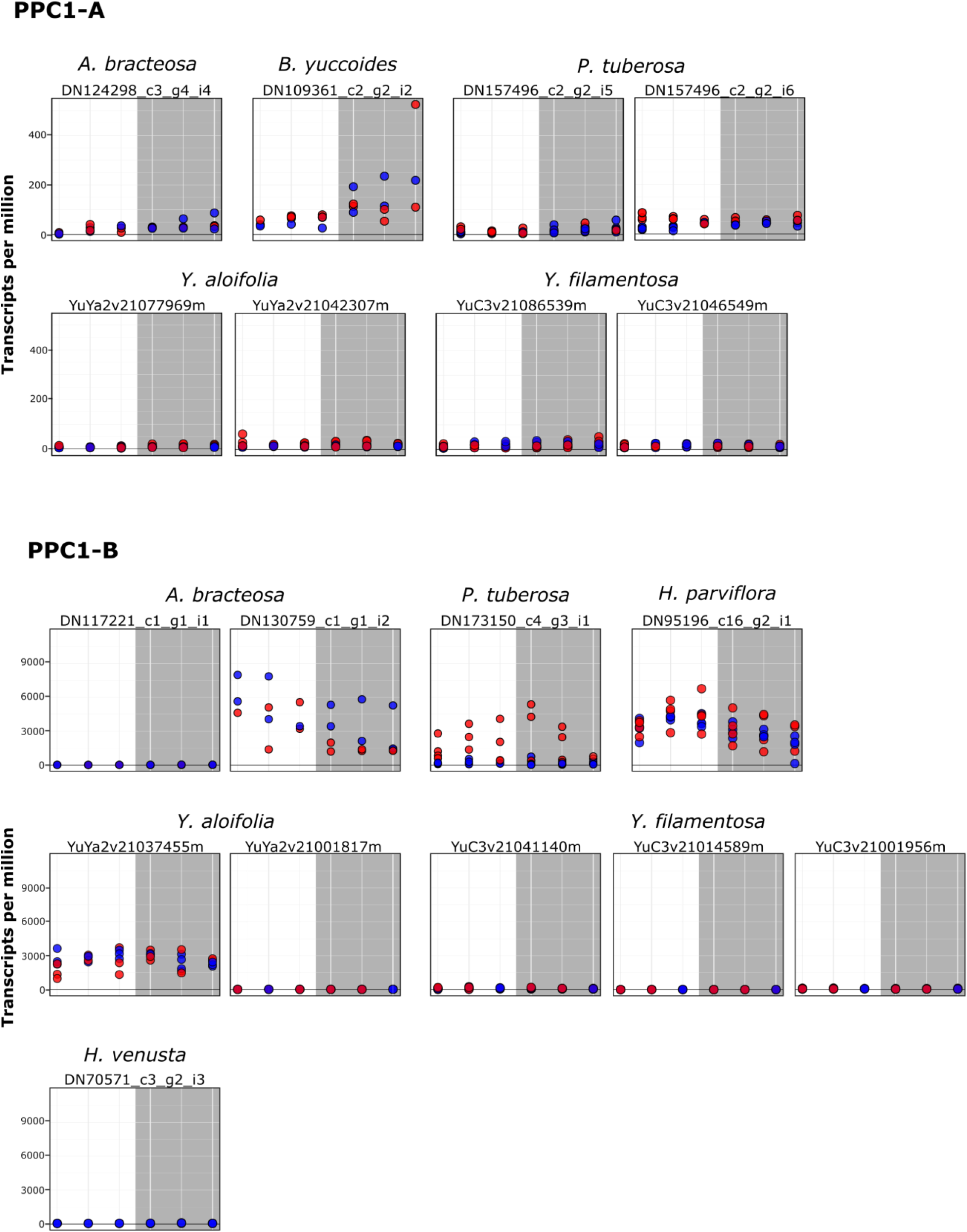
Expression (transcripts per million) of PPC1 transcripts from the core Agavoideae species, shown separately for the two clades A and B. Dots represent individual samples, with blue = well watered and red = drought stressed. Grey box indicates time points when the lights were off. Transcripts are only shown here if they passed length and percent identity filtering. *Hosta venusta* did not have drought-stressed samples taken for RNA-sequencing.

**Figure 6.**
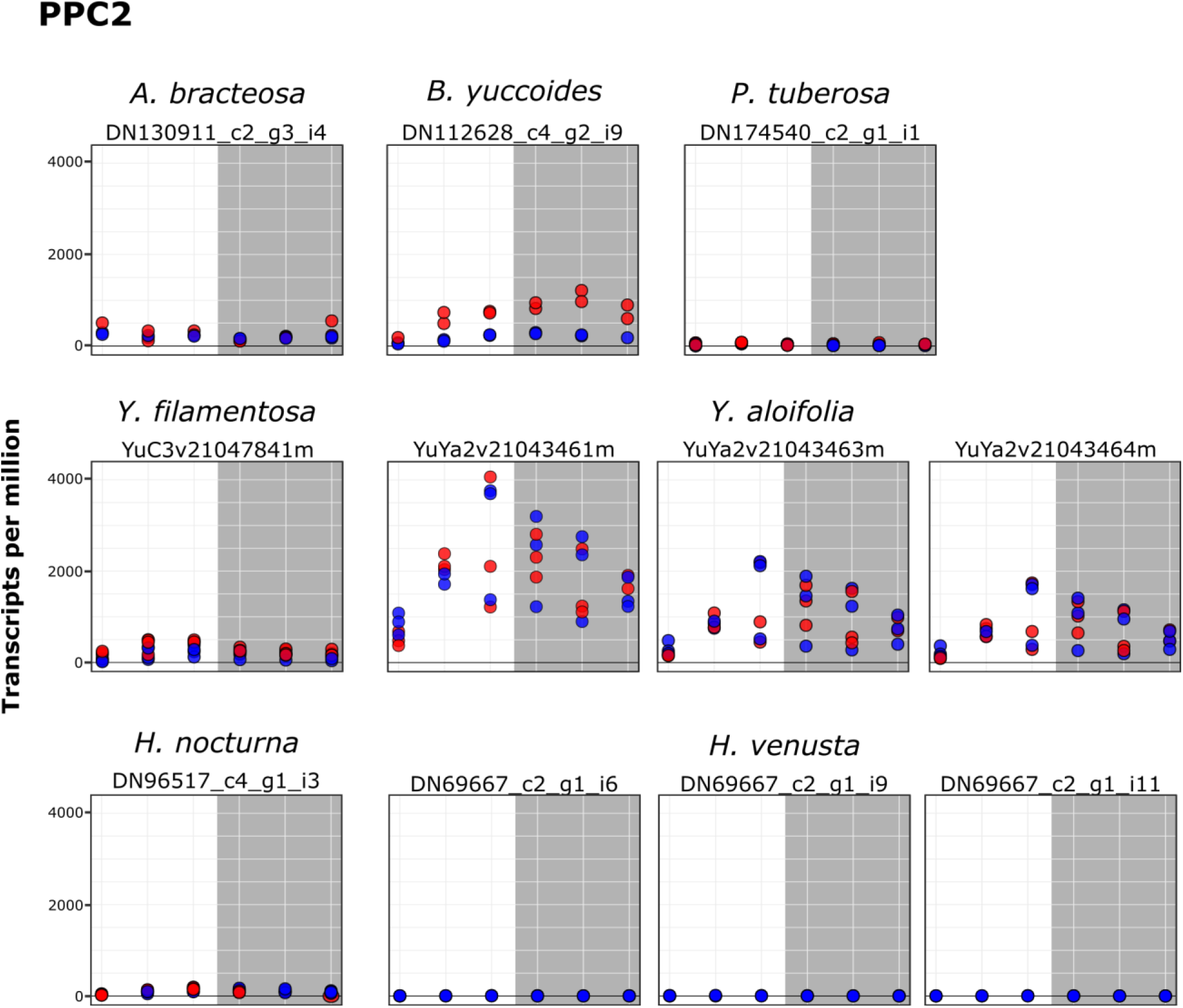
Expression (transcripts per million) of PPC2 transcripts from the core Agavoideae species. Dots represent individual samples, with blue = well-watered and red = drought stressed. Grey box indicates time points when the lights were off. Transcripts are only shown here if they passed length and percent identity filtering.

### Molecular evolution of PPC genes

Assessment of changes in the strength and mode of selection assessed by the branch model revealed a significant shift in ω for *PPC1-A*, but not *PPC1-B* or *PPC2*. PPC1-A had a reduced ω relative to the background rate, suggesting increased purifying selection consistent with this gene’s role in housekeeping pathways (Fig. 4, Table 1, Table 2). The sites model tests for positive selection were not significant for either *PPC1* or *PPC2* in the Agavoideae. However, *PPC1-B* had significant positive selection on some sites in the Agavoideae genes (Table 1), and Bayesian Empirical Bayes analysis revealed only one site under positive selection with a posterior probability > 95%: a transition from an alanine to an asparagine at position 591. *PPC1-B* also exhibited shifts on constraint in the clade-sites test, with the Agavoideae having a third class of sites with weaker purifying selection compared to the background rate (0.44 on the foreground, 0.21 on the background, proportion of sites = 0.27). *PPC2* likewise only had a significant rejection of the sites null model in favor of the alternative clade model, with a third class of sites that had an elevated ω relative to background (0.52 on foreground, 0.18 on background, proportion of sites = 0.36) (Table 2). Together these results suggest that specific amino acid residues in Agavoideae *PPC1-B* and *PPC2* genes may be evolving under relaxed or positive selection.

**Table 1.**
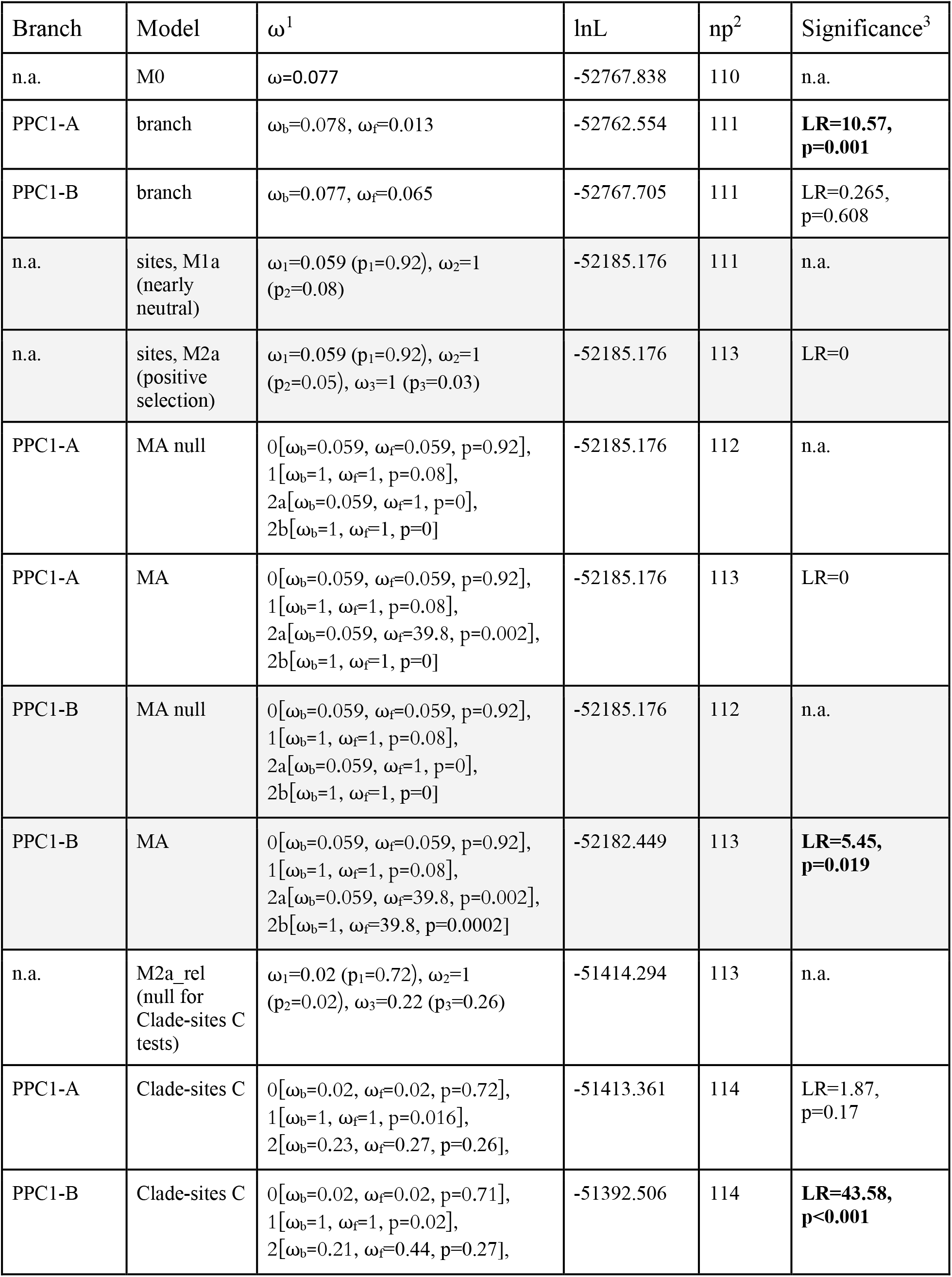
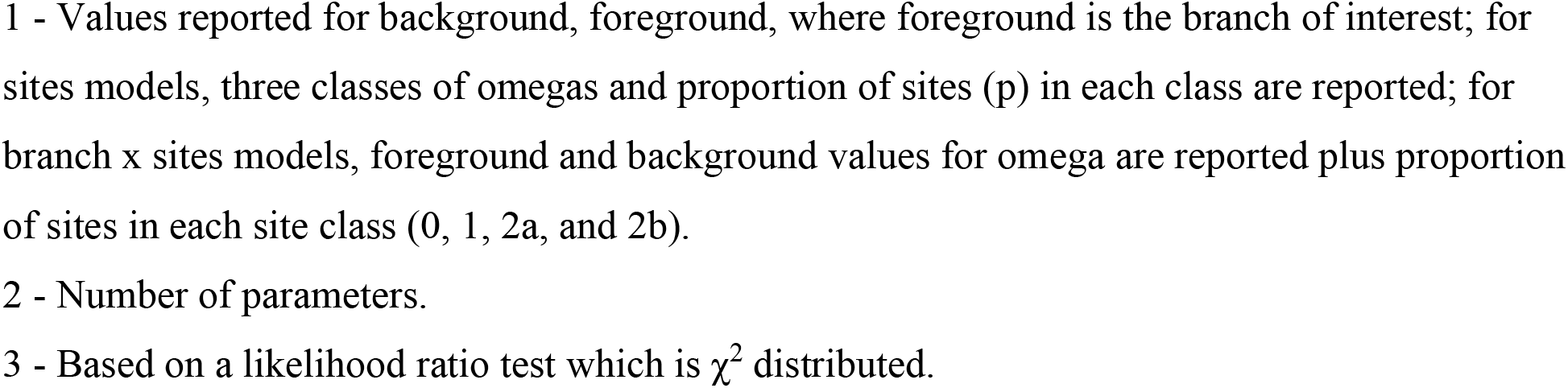
Results from tests of selection on PPC1.

| Branch | Model | $\omega^1$ | lnL | np<sup>2</sup> | Significance<sup>3</sup> |
| --- | --- | --- | --- | --- | --- |
| n.a. | M0 | $\omega=0.077$ | -52767.838 | 110 | n.a. |
| PPC1-A | branch | $\omega_b=0.078, \omega_f=0.013$ | -52762.554 | 111 | **LR=10.57,**<br>**p=0.001** |
| PPC1-B | branch | $\omega_b=0.077, \omega_f=0.065$ | -52767.705 | 111 | LR=0.265,<br>p=0.608 |
| n.a. | sites, M1a<br>(nearly<br>neutral) | $\omega_1=0.059$ ($p_1=0.92$), $\omega_2=1$<br>($p_2=0.08$) | -52185.176 | 111 | n.a. |
| n.a. | sites, M2a<br>(positive<br>selection) | $\omega_1=0.059$ ($p_1=0.92$), $\omega_2=1$<br>($p_2=0.05$), $\omega_3=1$ ($p_3=0.03$) | -52185.176 | 113 | LR=0 |
| PPC1-A | MA null | 0 [$\omega_b=0.059, \omega_f=0.059, p=0.92$],<br>1 [$\omega_b=1, \omega_f=1, p=0.08$],<br>2a [$\omega_b=0.059, \omega_f=1, p=0$],<br>2b [$\omega_b=1, \omega_f=1, p=0$] | -52185.176 | 112 | n.a. |
| PPC1-A | MA | 0 [$\omega_b=0.059, \omega_f=0.059, p=0.92$],<br>1 [$\omega_b=1, \omega_f=1, p=0.08$],<br>2a [$\omega_b=0.059, \omega_f=39.8, p=0.002$],<br>2b [$\omega_b=1, \omega_f=1, p=0$] | -52185.176 | 113 | LR=0 |
| PPC1-B | MA null | 0 [$\omega_b=0.059, \omega_f=0.059, p=0.92$],<br>1 [$\omega_b=1, \omega_f=1, p=0.08$],<br>2a [$\omega_b=0.059, \omega_f=1, p=0$],<br>2b [$\omega_b=1, \omega_f=1, p=0$] | -52185.176 | 112 | n.a. |
| PPC1-B | MA | 0 [$\omega_b=0.059, \omega_f=0.059, p=0.92$],<br>1 [$\omega_b=1, \omega_f=1, p=0.08$],<br>2a [$\omega_b=0.059, \omega_f=39.8, p=0.002$],<br>2b [$\omega_b=1, \omega_f=39.8, p=0.0002$] | -52182.449 | 113 | **LR=5.45,**<br>**p=0.019** |
| n.a. | M2a_rel<br>(null for<br>Clade-sites C<br>tests) | $\omega_1=0.02$ ($p_1=0.72$), $\omega_2=1$<br>($p_2=0.02$), $\omega_3=0.22$ ($p_3=0.26$) | -51414.294 | 113 | n.a. |
| PPC1-A | Clade-sites C | 0 [$\omega_b=0.02, \omega_f=0.02, p=0.72$],<br>1 [$\omega_b=1, \omega_f=1, p=0.016$],<br>2 [$\omega_b=0.23, \omega_f=0.27, p=0.26$], | -51413.361 | 114 | LR=1.87,<br>p=0.17 |
| PPC1-B | Clade-sites C | 0 [$\omega_b=0.02, \omega_f=0.02, p=0.71$],<br>1 [$\omega_b=1, \omega_f=1, p=0.02$],<br>2 [$\omega_b=0.21, \omega_f=0.44, p=0.27$], | -51392.506 | 114 | **LR=43.58,**<br>**p<0.001** |

1 - Values reported for background, foreground, where foreground is the branch of interest; for sites models, three classes of omegas and proportion of sites (p) in each class are reported; for branch x sites models, foreground and background values for omega are reported plus proportion of sites in each site class (0, 1, 2a, and 2b).
2 - Number of parameters.
3 - Based on a likelihood ratio test which is $\chi^2$ distributed.

**Table 2.** Results from tests of selection on PPC2.

| Model | $\omega^1$ | lnL | np<sup>2</sup> | Significance<sup>3</sup> |
| --- | --- | --- | --- | --- |
| M0 | $\omega=0.126$ | -27536.821 | 44 | n.a. |
| branch | $\omega_b=0.126, \omega_f=0.115$ | -27536.753 | 45 | LR=0.136, p=0.71 |
| sites, M1a<br>(nearly<br>neutral) | $\omega_1=0.07$ ($p_1=0.82$), $\omega_2=1$ ($p_2=0.18$) | -26777.829 | 45 | n.a. |
| sites, M2a<br>(positive<br>selection) | $\omega_1=0.07$ ($p_1=0.82$), $\omega_2=1$ ($p_2=0.09$), $\omega_3=1$ ($p_3=0.09$) | -26777.829 | 47 | LR=0 |
| MA null | 0 [$\omega_b=0.065, \omega_f=0.065, p=0.82$],<br>1 [$\omega_b=1, \omega_f=1, p=0.18$],<br>2a [$\omega_b=0.07, \omega_f=1, p=0$],<br>2b [$\omega_b=1, \omega_f=1, p=0$] | -26777.829 | 46 | n.a. |
| MA | 0 [$\omega_b=0.065, \omega_f=0.065, p=0.82$],<br>1 [$\omega_b=1, \omega_f=1, p=0.18$],<br>2a [$\omega_b=0.07, \omega_f=1, p=0$],<br>2b [$\omega_b=1, \omega_f=1, p=0$] | -26777.829 | 47 | LR=0 |
| M2a_rel<br>(null for<br>Clade-sites<br>C tests) | $\omega_1=0.02$ ($p_1=0.59$), $\omega_2=1$ ($p_2=0.09$), $\omega_3=0.23$ ($p_3=0.33$) | -26537.435 | 47 | n.a. |
| Clade-sites<br>C | 0 [$\omega_b=0.01, \omega_f=0.01, p=0.55$],<br>1 [$\omega_b=1, \omega_f=1, p=0.09$],<br>2 [$\omega_b=0.18, \omega_f=0.52, p=0.36$], | -26508.092 | 48 | **LR=58.68, p<0.001** |

1 - Values reported for background, foreground, where foreground is the branch of interest; for sites models, three classes of omegas and proportion of sites in each class are reported; for branch x sites models, foreground and background values for omega are reported plus proportion of sites in each site class (0, 1, 2a, and 2b).
2 - Number of parameters.
3 - Based on a likelihood ratio test which is $\chi^2$ distributed.

## Discussion

### Evolution of CAM in the Agavoideae

Previous work estimated three independent origins of CAM in the Agavoideae: one in the genus *Hesperaloe,* one in the genus *Yucca*, and one in *Agave s.l. (Heyduk et al., 2016b)*. However, this initial estimation was based on carbon isotope values, which cannot separate C_3_+CAM from C_3_ in the majority of cases (Winter et al., 2015). Detailed physiological measurements under both well-watered and drought-stressed conditions revealed that *Polianthes* has the ability to upregulate CAM under drought stress, though maintains low-level CAM even under well-watered conditions. *Beschorneria* has a very slight CAM increase at night, indicated by a small shift in titratable acidities and a decrease in nighttime respiration (Heyduk et al., 2018b). *Yucca* species are divided, in that nearly half are expected to use C_3_ and the other half are likely CAM; these inferences are based on carbon isotopes, and warrant more detailed physiological assessment beyond the two species included in this study and a handful of others (Smith et al., 1983; Heyduk et al., 2016a). The presence of CAM was confirmed in *Hesperaloe*, with both species in this study exhibiting strong CAM (Fig. 2). *Hosta* showed no evidence of CAM in our study, although it was not drought stressed. Based on gene expression patterns detected here, as well as anatomical traits and carbon isotope values (Heyduk et al., 2016b), we do not expect *Hosta* to be able to up-regulate CAM under drought stress. Together these separate physiological assessments across the Agavoideae confirm the presence of CAM in *Hesperaloe, Yucca,* and *Agave s.l.* and further our understanding of intermediate CAM species (e.g., *Polianthes* and *Beschorneria*).

### Conservation and novelty in gene expression

Across diverse plant species, roughly 20-60% of transcripts show some time-of-day differential expression (Covington et al., 2008; Hayes et al., 2010; Filichkin et al., 2011; Lai et al., 2020), however *Arabidopsis* has up to 89% of transcripts cycling under at least one experimental time course condition (Michael et al., 2008). In *Sedum album*, which has the ability to facultatively up-regulate CAM, there is a slight increase in the number of cycling transcripts when plants use CAM compared to C_3_ (35% vs. 41%, respectively) (Wai et al., 2019). The number of cycling transcripts in Agavoideae species varied, with *Hosta* having the fewest transcripts that were time-structured, and *Agave* having the greatest number. The number of time-structured transcripts did not cleanly correlate to the presence of CAM; for example, *Y. filamentosa* and *Y. aloifolia* had similar numbers of time-structured transcripts, despite differences in photosynthetic pathway. The lack of association between photosynthetic pathway and time-structured gene expression instead suggests ancestral gene networks in the genus *Yucca* were retained in both photosynthetic types, and is further supported by the high number of gene families that are time-structured in both species (n=639) (Fig. 3B). Whether or not the presence of time-structured gene expression networks in the ancestor of *Yucca* facilitated the evolution of CAM remains undetermined. Similarly, *Agave* and *Polianthes* both had a large number of transcripts with diel variation, despite *Polianthes* being only weakly, facultatively CAM. *Beschorneria*, which is sister to *Polianthes* and *Agave*, showed the smallest number of time-structured transcripts, though it is also the weakest CAM species measured across these species.

Very few gene families had time-structured expression across all CAM species (n=105 in all CAM, n=126 in strong CAM). Two key genes related to CAM — *PPC2* (though, notably, not *PPC1*) and *PPCK* — were time-structured in all species except *Hosta* (i.e., including the C_3_ *Y. filamentosa*). *PPCK* in particular has been shown to have direct and reciprocal clock connections; knock-downs of *PPCK* in *Kalanchoë fedtschenkoi* had significantly reduced CAM and the lack of circadian oscillation in *PPCK* perturbed oscillation patterns of core clock genes (Boxall et al., 2017). Knock-down of *PPC1* in *K. fedtschenkoi* also resulted in changes to the oscillation patterns and amplitude of clock genes, though notably a different set of core clock genes were affected by *PPC1* knockdowns relative to *PPCK (Boxall et al., 2020)*. The integration of the circadian clock and CAM pathway genes is clearly important for CAM physiology, though the presence and cycling of these genes does not, alone, lead to CAM ability - *Y. filamentosa* has cycling of these gene families (e.g., *PPC2*), but expression levels are either too low or transcripts are affected by other post-translational modifications to render them insufficient for CAM (Heyduk et al., 2019b). Moreover, we found a lack of shared gene families with shifts to time-structured expression across CAM species in the Agavoideae, and suggests three hypotheses: 1) the repeated evolution of CAM has involved lineage-specific changes to the molecular networks rather than parallelisms, 2) gene re-wiring happened in the ancestor of the Agavoideae and facilitated the repeated evolution of CAM, or 3) the overall scope of re-wiring of gene expression into the clock is limited for CAM.

Assessing the 24-hour time-structured variation of gene expression in CAM and C_3_ lineages has confirmed the important role of clock integration with CAM metabolic genes, but generally has not revealed any master regulator of CAM. In general the majority of studies highlight the conservation of circadian clock components and the timing of their expression, regardless of photosynthetic pathway (Moseley et al., 2018; Wai and VanBuren, 2018; Yin et al., 2018; Wai et al., 2019). Instead, researchers have focused on the few aspects of the clock that are different between C_3_ and CAM comparisons, but it’s worth noting that those comparisons are often between distantly related species, and it’s unclear whether these changes are therefore related to the evolution of CAM or simply stochastic changes in circadian network coordination. For example, *PRR9* in *Opuntia* (CAM) was shown to have a change in phase compared to the *Arabidopsis* ortholog (Mallona et al., 2011), comparisons between *Kalanchoe* and *Arabidopsis* showed phase shifts in a number of evening elements, including *ELF3/4* and *LUX* (Moseley et al., 2018), and *Agave* had shifted expression of RVE, a clock output gene, relative to *Arabidopsis* (Yin et al., 2018). In the Agavoideae, the majority of circadian gene families have shared patterns of time-structured expression across all eight species. Of those gene families that had a significant species effect in the phase of expression, few had extreme phase shifts or showed consistent C_3_ vs. CAM differences. Our findings, together with those of other studies assessing core circadian regulators in CAM lineages, point to an overall conservation of the circadian clock, even in plants with a strong CAM physiology (Boxall et al., 2020). However, it’s worth noting that the majority of comparative transcriptomics studies in CAM, including this one, assess temporal variation in expression over a single day-night period, making it difficult to pinpoint which genes are responsible for clock inputs into the CAM pathway, and which are downstream targets. Many studies still rely on distant outgroups for comparison (typically *Arabidopsis*), and thus continue to confound changes associated with CAM to those that arise simply due to evolutionary divergence. Future work on the nature of gene expression and evolution in CAM species should endeavor to use free-running conditions to better assess the roles of the circadian clock in CAM species, and should carefully select comparison species to minimize evolutionary distance. Regardless, it seems unlikely that large perturbations to the circadian clock are required for the evolution of CAM from a C_3_ ancestor; instead, changes to promoter sequences and regulatory regions of genes contributing to CAM may have a larger role to play in altering the timing and magnitude of their expression.

Unlike the relatively conserved number and type of genes that exhibited time-structured variation in gene expression, the response to drought was highly lineage specific. A large proportion of gene families were uniquely differentially expressed in a singular species. While the present study includes comparisons across three separate experiments (Heyduk et al., 2018b; Heyduk et al., 2019b), even species drought-stressed in the same experiment show vastly different responses to drought. *Agave*’s strong differential regulation to drought is surprising, given its constitutive CAM physiology is thought to buffer against the effects of drought stress. Indeed, the majority of CAM species studied here were affected by drought: *Y. aloifolia, A. bracteosa,* and *Manfreda sp.* all exhibited decreases in titratable leaf acidity and, in *Yucca* and *Manfreda*, drops in nocturnal CO_2_ assimilation. The effects of drought stress on CAM physiology are vastly understudied, although work has been done in facultative CAM species (Cushman et al., 2008; Wai et al., 2019; Heyduk et al., 2020). Both the physiological and gene expression data presented here suggests full CAM species are not immune to effects of drought, and indeed exhibit strong physiological and transcriptional responses. Finally, for all species studied here, the majority of drought-responsive genes were also time-structured; in other words, constitutively expressed genes were infrequently affected by drought stress.

### Gene recruitment for CAM photosynthesis

In all published instances of C_4_ or CAM evolution, the PPC gene copy that gets recruited is from a gene family known as the “plant” PPCs - or PPC1. PPC1 is used by all plants for the replenishment of intermediates in the TCA cycle, and a singular copy typically gets re-wired for C_4_ or CAM (Heyduk et al., 2019a). The clear CAM-like expression of *PPC2* in *Yucca aloifolia* and, to a lesser extent, *Beschorneria yuccoides,* suggests that both of these species have recruited PPC2 as an alternate carboxylating enzyme for CAM. *PPC1* is still expressed in both of these species, and supports previous work that suggests PPC2 forms a hetero-octamer with PPC1 (O’Leary et al., 2011), though this remains to be tested in the Agavoideae. While we cannot say for certain PPC2 protein is produced, it seems unlikely that the transcripts would be expressed so highly (>1000 TPM) in *Yucca aloifolia* with no functional consequence. Moreover, expression of *PPC2* in *Yucca aloifolia* peaks right before the onset of the night period, consistent with expression patterns of PPCs in other Agavoideae in this study, as well as expression profiles of PPC in other lineages (Ming et al., 2015; Abraham et al., 2016; Yang et al., 2017; Heyduk et al., 2018a; Wai et al., 2019). The overall carboxylase activity of PPC2 in the Agavoideae remains to be studied, but could lend further clues as to how this atypical gene copy was recruited into the CAM pathway.

Unlike C_4_ PPCs, where convergent amino acid substitutions seem key to the recruitment of *PPC1* gene copies into the C_4_ pathway (Christin et al., 2007; Rosnow et al., 2014; Goolsby et al., 2018), evidence for convergent evolution at the molecular level in CAM is lacking. A comparison of PPC sequences between *Kalanchoe* and *Phalaenopsis* did reveal a shared amino acid change from R/H/K to D and was shown to significantly increase the activity of PPC (Yang et al., 2017). However, this amino acid substitution is not ubiquitous in CAM species; it is absent from *Ananas (Yang et al., 2017)* and all members of the Agavoideae examined here (see github repository for fasta files), suggesting that either the shared mutation is due to homoplasy, or may be convergent but not essential for CAM. Our results further suggest that overall *PPC* genes are conserved, even when they are being recruited into the CAM pathway (Tables 1 and 2). In general, the lability of CAM as a phenotype, as well as the wide diversity of lineages in which it evolves, seems to allow variable pathways to organize the genetic requirements, including which major copy of the main carboxylating enzyme, PPC, is recruited. Increasing number of CAM lineages studied physiologically and genomically will allow us to determine whether novel mechanisms of evolving CAM — like the recruitment of *PPC2* in the Agavoideae — are indeed rare, or more common across green plants.

## Conclusions

By comparing RNA-seq data across closely related species that span multiple origins of CAM, we have shown that the majority of gene families have diurnal variation in gene expression, regardless of photosynthetic status. In particular, core circadian clock genes are similarly expressed across all the species examined here. In contrast, drought response was highly lineage specific, and suggests lineages have fine-tuned or independently evolved their drought response gene networks. While historically CAM in the Agavoideae has been thought to be the result of three independent origins, we cannot rule out a single origin of CAM with subsequent reversals to C_3_. However, reversals to C_3_ from CAM appear to be rare in angiosperms, and the recruitment of *PPC2* for CAM function in *Yucca* (and to a lesser extent in *Beschorneria*) supports the inference of independent origins of CAM in the Agavoideae and furthers the idea that the evolutionary routes to CAM are remarkably variable.

## Materials and methods

### Plant growth and physiological sampling

Plants of *Hesperaloe parviflora* (accession: PARL 436) and *Hesperaloe nocturna* (accession: PARL 435) were grown from seed acquired in 2014 from the USDA Germplasm Resources Information Network (GRIN). *Hesperaloe* plants were kept in the University of Georgia (UGA) Plant Biology greenhouses with once weekly watering. *Hosta* plants were purchased for New Hampshire Hostas (https://www.nhhostas.com/) in January 2018 and kept on a misting bench at the same greenhouses until experimentation began in March 2018. Replicates of each species (n=4, 4, and 6 for *H. parviflora, H. nocturna,* and *H. venusta*, respectively) were placed into a walk-in Conviron growth chamber, with day length set to 12 hours (lights on at 7 a.m.), day/night temperatures 30/17°C, humidity at 30%, and maximum PAR (about 400 μmol m^−2^ s^−1^ at plant level).

Plants were acclimated in the growth chamber for four days prior to sampling and watered to saturation daily. On day 1, plants were sampled every two hours, beginning 1 hour after the lights turned on (7 a.m.), for gas exchange using a LI-COR 6400XT. Due to the small size of the plants, only two replicates of *Hosta* had LI-COR measurements taken; one replicate of *Hesperaloe nocturna* was not measured due to an ant infestation in the pot. After day 1, water was withheld for five days in all plants with the exception of *Hosta*, which were all removed from the experiment at this point. On day 7, all remaining plants’ water status had dropped 8% soil water content, and plants were measured again for gas exchange. After day 7, plants were re-watered, and one more day of gas exchange sampling was conducted on day 9. Triplicate leaf tissue samples per plant were collected for titratable acidity measurements 2 hours before lights turned on (pre-dawn sample) and 2 hours before lights turned off (pre-dusk sample). Samples for leaf titrations were immediately flash frozen and stored at −80 °C until measurement.

Leaf acid titrations were conducted as in (Heyduk et al., 2018b); briefly, frozen leaf disks were quickly weighed and placed into 60 mL of 20% EtOH. Samples were boiled until volume reduced to half, then 30 mL of diH_2_O was added. Samples were reduced to half again and a final volume of 30 mL of diH2O was added. Samples were allowed to cool then titrated to pH 7.0 using 0.002 M NaOH. Total μmoles H+ per gram of frozen mass was calculated as (mL NaOH × 0.002 M)/g. Pre-dusk values were subtracted from pre-dawn to get the change, or ΔH+, per replicate. All statistical analyses were conducted in R v 3.5.0 (R Core Team, 2019).

### RNAsequencing and assembly

Tissue for RNA-sequencing was collected every four hours from each of the three species, from four replicate plants per species. For *H. nocturna* and *H. parviflora*, samples were collected from both well-watered and drought-stressed plants (days 1 and 7). For *Hosta venusta,* only well-watered samples were collected. Tissue was flash frozen in N2, then stored at −80 °C. RNA was isolated using a QIAGEN RNeasy Plant Kit, purified with Ambion Turbo DNAse, and quantified via a nanodrop and Agilent Bioanalyzer v2100. RNA-sequencing libraries were constructed with a KAPA mRNA Stranded kit at half reaction volume and barcoded separately using dual barcodes (Glenn et al., 2019). Library concentrations were measured via quantitative PCR, pooled in sets of 28-29 libraries, and sequenced with PE 75bp reads on an Illumina NextSeq at the Georgia Genomics and Bioinformatics Core at the University of Georgia. Raw reads from sequencing *Hesperaloe* and *Hosta* species are available on NCBI’s SRA, under BioProject PRJNA755802.

Raw reads were processed with Trimmomatic v 0.36 (Bolger et al., 2014) and paired reads were assembled *de novo* for each of the three species (*Hesperaloe parviflora, H. nocturna,* and *Hosta venusta*) in Trinity v. 2.5.1 (Grabherr et al., 2011). Reads were initially mapped to the entire Trinity-assembled transcriptome for each species with Bowtie v2.0 (Langmead and Salzberg, 2012). Trinity “isoforms” that had less than 2 transcripts mapped per million (TPM) abundance or constituted less than 20% of total component expression were removed. Transcriptome assemblies of sister species, including *Agave bracteosa, Polianthes tuberosa,* and *Beschorneria yuccoides (Heyduk et al., 2018b)* had already been filtered by the same thresholds as above. All six filtered assemblies had open reading frames (ORFs) predicted by Transdecoder v. 2.1 (Grabherr et al., 2011) using both LongOrfs and Predict functions and keeping only the best scoring ORF per transcript.

To sort the predicted Transdecoder sequences into gene families, we generated orthogroups circumscribed from nine reference genomes downloaded from Phytozome (Goodstein et al., 2012), with a particular focus on monocots. Translated primary transcript sequences were downloaded for *Acorus americanus* v1.1 (DOE-JGI, http://phytozome-next.jgi.doe.gov/), *Arabidopsis thaliana* v. Araport11 (Cheng et al., 2017), *Asparagus officinalis* v.1.1 (Harkess et al., 2017), *Ananas comosus* v.3 (Ming et al., 2015), *Amborella trichopoda* v.1 (Amborella Genome Project, 2013), *Brachypodium distachyon* v.3.1 (International Brachypodium Initiative, 2010), *Dioscorea alata* v2.1 (DOE-JGI), *Musa acuminata* v.1 (D’Hont et al., 2012), *Oryza sativa* v.7 (Ouyang et al., 2007), *Sorghum bicolor* v.3.1.1 (McCormick et al., 2018), and *Setaria italica* v.2.2 (Bennetzen et al., 2012) from Phytozome. Translated coding sequences from these genomes were clustered using OrthoFinder v.2.2.7 (Emms and Kelly, 2019). In addition to the above published genomes, preliminary draft genome data (primary translated transcripts) for *Yucca aloifolia* and *Yucca filamentosa* were secondarily added to the orthogroup analyses using the −b flag of OrthoFinder. Finally, Transdecoder translated coding sequences for the Agavoideae species were then added to the orthogroup circumscription again using the −b flag.

### Expression analysis

Reads were remapped using Kallisto (Bray et al., 2016) onto the filtered transcriptomes (iso_pct > 20, TPM > 2, Transdecoder best scoring ORF) for the *de novo* assemblies of *Hesperaloe* and *Hosta*. For the two *Yucca* genomes, existing RNA-seq reads (from (Heyduk et al., 2019b) were mapped onto the annotated primary transcripts using Kallisto. Because previously published expression analysis of *Agave, Beschorneria,* and *Polianthes* was done on transcriptomes filtered the same way (iso_pct>20, TPM>2), expression data in the form of read counts and TPM values for genes were used as previously published. Count and TPM matrices for all taxa analyzed here, as well as orthogroup annotations, are available on github (www.github.com/kheyduk/AgavoideaeComparative).

Read counts for the two *Hesperaloe* species, *Hosta,* and the two *Yucca* species were imported into R for initial outlier filtering in EdgeR (Robinson et al., 2010) and subsequent time-structured expression analysis in maSigPro (Conesa et al., 2006; Nueda et al., 2014). The latter program fits read count data to regressions, taking into account treatments (well-watered and drought stress), and asks whether a polynomial regression of degree *n* (chosen to be 5, or one less the number of timepoints) is a better fit to each gene than a straight line. Genes with expression patterns across time can be best explained by a polynomial regression are hereafter referred to as “time-structured.” While read counts are required for the maSigPro analysis, all comparative expression plots presented use transcripts mapped per million (TPM) normalized expression.

### Circadian gene expression

Previous studies in CAM have shown that a few circadian regulators get re-wired in the evolution of CAM (Moseley et al., 2018; Wai et al., 2019). To determine whether these patterns hold in more closely related C_3_ and CAM species, expression of circadian clock genes was compared between members of the Agavoideae. From the list of genes that had significantly time-structured expression from maSigPro for each species, we assessed gene family presence/absence data from the OrthoFinder gene circumscriptions. Shared gene family presence in the time-structured expression was assessed using the UpSetR package (Conway et al., 2017) in R 4.0.4 (R Core Team, 2013). A curated list of *Arabidopsis thaliana* circadian genes was used to examine the extent to which time-structured expression of circadian genes was shared across all eight species. Finally, for circadian-annotated genes, we employed JTK_CYCLE (Hughes et al., 2010) and Lomb-Scargle (Glynn et al., 2005) methods implemented in MetaCycle (Wu et al., 2016) to obtain period, lag, and amplitude for genes with a period expression pattern. Cycling patterns for *Agave* and *Beschorneria* were excluded from further analysis, as their resolution (number of replicates and time points) was lower than other species due to dropped libraries (Heyduk et al., 2018b). We then used ANOVA to assess whether there were differences in average phase across OrthoFinder gene families between CAM and C_3_ species, as well as between *Hosta* and the other Agavoideae species. P-values were corrected for multiple testing using Benjamini-Hochberg.

### PPC evolution

The *PPC1* and *PPC2* gene families were identified in the OrthoFinder-circumscribed orthogroups by searching for annotated *Arabidopsis PPC1* and *PPC2* copies. Both orthogroups were manually inspected for completeness by checking if all known genes from sequenced and annotated genomes were properly sorted into those two orthogroups. Only sequences that were at least 50% the length of the longest sequence (based on coding sequence) were retained and aligned. Sequences from the *de novo* transcriptomes were collapsed (using the longest as the representative) within a species if they were >96.32% and >99.71% identical for *PPC1* and *PPC2*, respectively. To get in-frame coding sequence alignment, each orthogroup protein and CDS output from Transdecoder were used to align the coding sequences using PAL2NAL (Suyama et al., 2006). Phylogenetic trees for *PPC1* and *PPC2* were estimated on the in-frame coding sequence alignments using IQtree v2.0 (Nguyen et al., 2015; Minh et al., 2020) and 1000 rapid bootstrap replicates, using built-in ModelFinder to determine the best substitution model. The resulting tree for each gene family, along with the in-frame coding sequence alignment, were used to estimate shifts in molecular evolution using codeml in PAML (Yang, 2007). Specifically, we tested branch, sites, and branch(clade)-sites models. We compared branch models to a null M0 model with a single ω value, the M2a sites model (positive selection) to the null M1a (nearly neutral)(Wong et al., 2004), the branch-sites model A to the null (fixed_omega=1, omega=1), and the clade C model to M2a_rel. For branch, branch-sites, and clade models, we labeled the two *PPC1* Agavoideae lineages and estimated ω separately; for *PPC2*, we labeled the single Agavoideae stem branch. Due to low phylogenetic resolution within the Agavoideae, specific tests for independent CAM origins were not feasible. Fasta files of multispecies alignments and newick gene trees are available at www.github.com/kheyduk/AgavoideaeCAM.

## Supplemental Materials

**Supplemental Table S1** – Raw Li-COR data for *Hesperaloe parviflora, Hesperaloe nocturna*, and *Hosta venusta*.

**Supplemental Table S2** – Titratable acidity measurements for *Hesperaloe parviflora, Hesperaloe nocturna*, and *Hosta venusta*.

**Supplemental Table S3** – Gene families with time-structured expression across all eight species.

**Supplemental Table S4** – Gene families with time-structured expression in all species but *Hosta venusta*.

**Supplemental Table S5** – Analysis of circadian annotated genes in Metacycle.

**Supplemental Table S6** – Abbreviated ANOVA results of testing effect of species on phase of gene expression.

## Acknowledgements

The authors would like to acknowledge the UGA Plant Biology greenhouse staff for the help in maintaining plants, the Georgia Advanced Computing Resources Center, and Amanda L. Cummings and Richard Field for assistance with greenhouse and lab work. We thank the Department of Energy Joint Genome Institute and collaborators for access to pre-publication data for *Acorus americanus* v1.1, *Dioscorea alata* v2.1, *Yucca aloifolia* and *Yucca filamentosa*.

